# Time-Resolved Single-Molecule FRET Reveals Length-Dependent Nucleosome Decompaction by Poly(ADP-ribose)

**DOI:** 10.64898/2026.02.25.707951

**Authors:** Tianjin Yang, Soundhararajan Gopi, Louise Pinet, Sebastian Simoni, Ralph Imhof, Daniel Nettels, Matthias Altmeyer, Robert B. Best, Benjamin Schuler

## Abstract

Highly charged chains of poly(ADP-ribose) (PAR) are synthesized in the cell as part of their central role in DNA damage response. However, the effects of PAR on nucleosome structure and dynamics remain incompletely understood. Here we combine droplet-based microfluidic mixing with single-molecule Förster resonance energy transfer spectroscopy to resolve the kinetics of PAR-induced nucleosome decompaction in non-equilibrium measurements with millisecond time resolution. This approach avoids surface-adhesion and enables the tether-free observation of nucleosome remodeling. We find that PAR triggers nucleosome decompaction via a length-dependent kinetic threshold: Chains with less than ten ADP-ribose units act slowly and weakly, whereas longer PAR polymers induce efficient and rapid nucleosome opening. The extent and reversibility of decompaction further depend on PAR concentration and ionic strength, reflecting a mechanism dominated by electrostatic interactions. Enzymatic PAR digestion demonstrates that PAR can promote both reversible linker DNA opening and irreversible nucleosome disassembly. Coarse-grained molecular simulations suggest that these effects arise from a competition between PAR and DNA for histone tail binding. Altogether, our results establish PAR length as a key factor controlling chromatin accessibility during DNA repair and highlight droplet-based microfluidics as a powerful platform for studying such biomolecular interactions.

## Introduction

Nucleosomes are the elementary packaging units of cellular DNA. The nucleosome is stabilized by electrostatic interactions between the negatively charged DNA and the positively charged histones^1, 2^. The intrinsically disordered tails of histones play essential roles in chromatin organization and gene regulation by modulating DNA packaging^2, 3^, nucleosome stability^2, 4–6^, and interactions with regulatory proteins^7^, serving as key sites for post-translational modifications (PTMs)^3, 8^. Among these PTMs, poly(ADP-ribosyl)ation (PARylation) of specific amino acids on histone tails is critical for DNA repair^9–13^. Single-strand breaks and double-strand breaks are detected by poly(ADP-ribose) polymerase 1 (PARP1)^14–17^. Upon binding to DNA breaks, PARP1 becomes activated and rapidly PARylates itself and histone octamers, generating poly(ADP-ribose) (PAR) - a negatively charged, nucleic acid-like polymer of variable length synthesized from NAD^+18, 19^. Recent results show that free PAR can also be synthesized de novo by PARP1, both biochemically and intracellularly in response to DNA damage and represents about 20–35% of the total PAR isolated from cells^20^. PAR acts as a scaffold, recruiting DNA repair proteins and facilitating the formation of condensates at damage sites^21–25^. In addition, some studies suggest that PAR alters the mobility of positively charged histone tails and their interactions with DNA within nucleosomes^26–32^, decompacting them and exposing damaged sites for repair. Despite these advances, the structural consequences of PAR interacting with nucleosomes remain poorly defined. In particular, which conformational changes of nucleosomes are triggered by PAR during DNA repair — partial linker opening or more extensive DNA unwrapping? How do the properties of PAR, especially its degree of polymerization, influence its interactions with the nucleosome?

In this study, we investigated the effect of PAR on the structural dynamics of nucleosomes by employing single-molecule fluorescence spectroscopy in combination with a droplet-based microfluidic mixer^33^ that prevents nucleosome surface adhesion, enabling direct, tether-free, and quantitative observation of PAR-induced nucleosome conformational changes under non-equilibrium conditions — a challenge for conventional methods. We find that PAR-induced nucleosome decompaction exhibits a threshold response whose rates and extent depend on PAR chain length, concentration, and salt concentration. Combining single-molecule experiments with coarse-grained simulations, we suggest that nucleosome decompaction is characterized by the loss of H3 and H2A C-terminal tail interactions with linker DNA due to the competition of histone tails with PAR. Our findings suggest PAR to be a dynamic regulator of nucleosome architecture, with length-dependent kinetics and salt sensitivity shaping its biological functions in DNA damage response and chromatin stability.

## Results

### Microfluidic single-molecule FRET reveals kinetics of PAR-induced nucleosome decompaction

To elucidate the interaction between poly(ADP-ribose) (PAR) and nucleosomes, as well as the process of PAR-induced nucleosome decompaction, we investigated the conformational distributions of nucleosomes and the kinetics of PAR-induced nucleosome decompaction under non-equilibrium conditions. This approach allows us to capture the rapid decompaction kinetics, which is much faster than the timescales accessible by manual mixing, and to observe even potentially irreversible processes^34–36^, such as histone dissociation from the nucleosome, or complete dissociation of DNA from the histone octamer. We reconstituted nucleosomes with 197 bp Widom DNA^37^ and labeled the termini of the linker DNA arms with Alexa 488 and Alexa 594 as FRET donor and acceptor dyes, respectively (**Figure 1a**, see Methods for details). In equilibrium experiments on freely diffusing molecules, the reconstituted nucleosomes exhibited two distinct populations: a major compact state with a high mean FRET efficiency (⟨E⟩ ≈ 0.36) and a minor population with low mean FRET efficiency (⟨*E*⟩ ≈ 0.06) (**Figure 1b**). Donor-only contributions to the signal from molecules lacking an active acceptor dye were eliminated by pulsed interleaved excitation^38^. The population at low ⟨*E*⟩ may thus be comprised of partially unwrapped nucleosomes and/or fully dissociated DNA according to the small population of free DNA detected electrophoretically in the sample (**Figure S1a right**) and the similarity to the low transfer efficiency observed for 197 bp Widom DNA without histones (**Figure 1d**).

**Figure 1:**
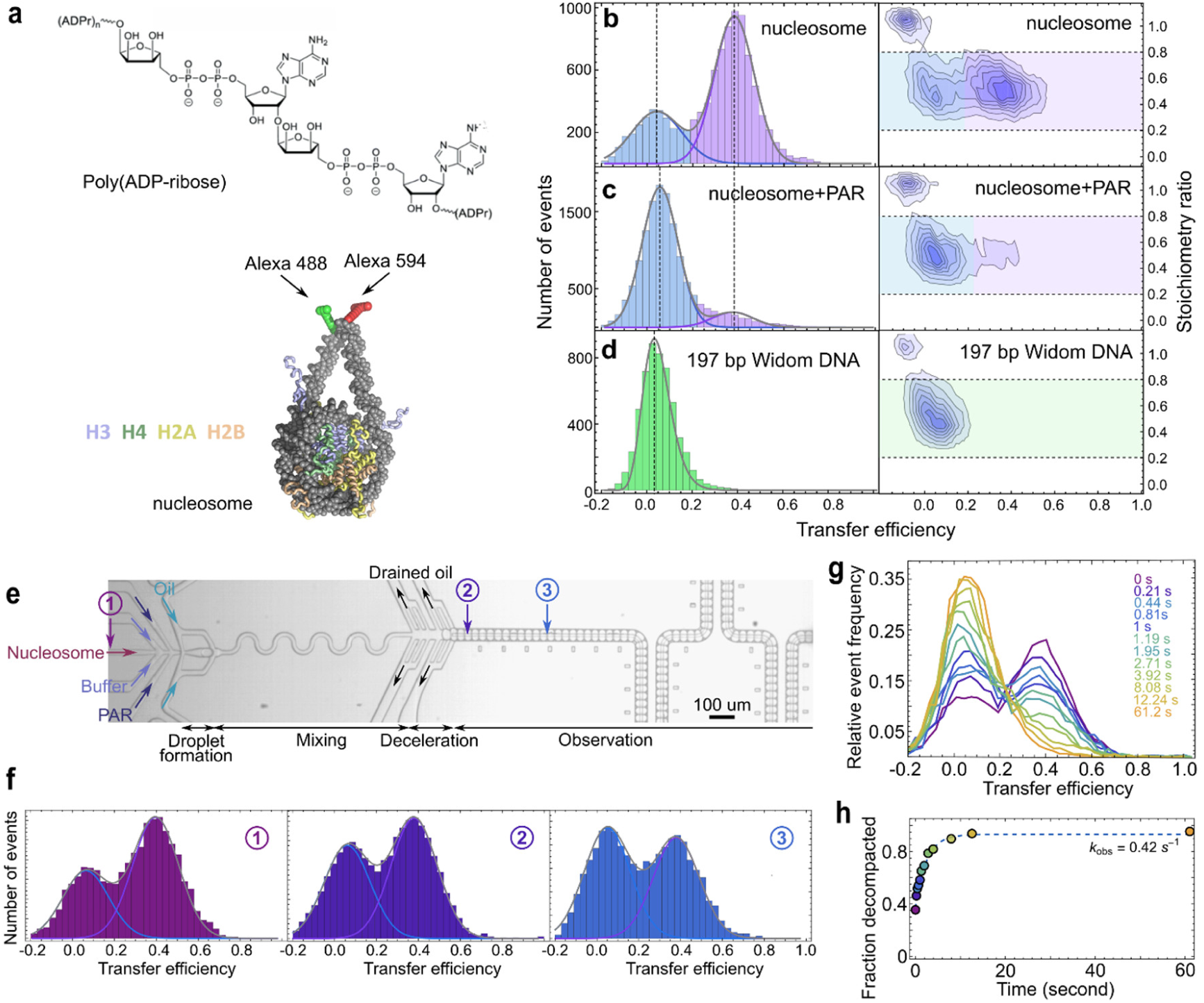
A droplet-based microfluidic system for single-molecule FRET reveals the kinetics of PAR-induced nucleosome decompaction. **a**, Illustration of poly(ADP-ribose) (PAR), and nucleosome labeled with FRET fluorophores Alexa 488 and Alexa 594 at the termini of the linker DNA arms. **b-d**, Histograms of transfer efficiency (left) and transfer efficiency vs stoichiometry ratio (right) of reconstituted nucleosomes (b), nucleosomes bound to PAR28-30 at equilibrium (c), and 197 bp Widom DNA without core histones (d) at an ionic strength of 300 mM. The reconstituted nucleosomes exhibit a major compact state (purple) with high mean FRET efficiency (⟨*E*⟩ ≈ 0.36, dashed line) and a minor population (blue) with low mean FRET efficiency (⟨*E*⟩ ≈ 0.06, dashed line). Upon interaction with PAR28-30 at equilibrium, the majority of the population at high ⟨*E*⟩ (purple) shifts to low ⟨*E*⟩ (blue). 197 bp Widom DNA without core histones (green) exhibits a mean FRET efficiency of ⟨*E*⟩ ≈ 0.035 (dashed line). The donor-only population with a stoichiometry ratio *S* of ∼1 was removed from the transfer efficiency histograms with a stoichiometry selection 0.2 < *S* < 0.8 (horizontal dashed lines). **e**, Widefield microscopy image of droplet formation, mixing, deceleration, and part of the inlets and observation channel of the droplet-based microfluidic mixer. Measurements were performed at various positions in the nucleosome inlet and along the observation channel. **f**, Examples of FRET efficiency histograms acquired at positions indicated with ①, ②, and ③. The peak at 〈*E*〉 ≈ 0.36 (purple fit line) corresponds to compact nucleosomes; the peak at〈*E*〉 ≈ 0.06 (blue fit line) corresponds to decompacted nucleosomes at times 210 ms (position ②) and 810 ms (position ③) after the start of the reaction (ionic strength 300 mM). The measurement at position ① was used for quantifying the initial population of folded nucleosomes. **g**, Normalized transfer efficiency histograms measured at different positions along the observation channel after mixing in droplets containing 100 pM nucleosome and 240 nM PAR28-30, corresponding to different times after the start of the reaction (ionic strength 300 mM). **h**, The kinetic data were fitted with a two-state model assuming a pseudo-first-order reaction (dashed line), yielding an observed decompaction rate of *k*_obs_ = 0.42 ± 0.08 s^−1^.

In prior kinetics measurements, we tested nucleosome stability at various KCl concentrations by measuring FRET population changes over time in free-diffusion experiments. We observed that the nucleosomes’ mean FRET efficiency, ⟨*E*⟩, ranged from a single peak of 0.089 ± 0.002 at low ionic strength to two peaks at higher ionic strengths (**Figure S2a**). The nucleosomes remained compact for at least five hours at an ionic strength of 300 mM or lower, four hours at an ionic strength of 390 mM, and three hours at an ionic strength of 495 mM (**Figure S2b**). The following measurements were completed within the time during which the nucleosomes remained stable.

To be able to probe the effect of PAR on the nucleosome, PAR was synthesized via PARP1-catalyzed NAD^+^ cleavage into nicotinamide and conjugated ADP-ribose units and separated by PAR chain length using strong anion exchange high-performance liquid chromatography (HPLC), and the degrees of polymerization were quantified by mass spectrometry (**Figure S3**). Addition of PAR to nucleosomes results in decompaction of the nucleosome (Figure 1c, addition of PAR with 28-30 units, PAR28-30). However, the process is too rapid to be time-resolved by manual mixing. We thus probed nucleosome decompaction kinetics caused by PAR using a droplet-based microfluidic mixer coupled with confocal single-molecule FRET spectroscopy^33^. The nucleosome and PAR solutions were delivered into the mixer and separated by a buffer stream prior to droplet formation to prevent premature reaction initiation. The PAR and nucleosome solutions were rapidly mixed in water-in-oil droplets and decelerated for single-molecule detection (Figure 1e). By encapsulating the nucleosomes in droplets, we overcome a key limitation of conventional laminar-flow microfluidic mixers^39–45^, where the highly positively charged histones tend to adhere strongly to channel surfaces, preventing kinetic measurements. The droplet-based microfluidic mixer instead enables tether-free, quantitative observation of PAR-induced nucleosome conformational changes under non-equilibrium conditions.

Nucleosome populations prior to mixing were measured at the device inlet (**Figure 1e, f**). Compared to free-diffusion measurements, the compact nucleosome population was slightly lower in the inlet of the mixing device but retained the same ⟨*E*⟩ (**Figure 1b**), indicating minor sample loss of the compact nucleosome at the inlet without conformational perturbation. Decompaction kinetics were recorded by placing the confocal volume at different positions along the observation channel (**Figure 1e, f**) corresponding to reaction times from several milliseconds to ∼60 seconds, with a droplet velocity of 0.6 mm/s. Upon mixing PAR with the labeled nucleosomes, the population with higher FRET efficiency at ⟨*E*⟩ = 0.36 ± 0.004 decreases, with a concomitant increase of the population at ⟨*E*⟩ = 0.053 ± 0.002, reflecting nucleosome decompaction (**Figure 1g**; uncertainties report experimental precision based on multiple measurements). The fraction of decompacted nucleosomes at each time was quantified and the results fitted to a two-state model under the assumption of pseudo-first-order reaction conditions to obtain the decompaction rate (**Figure 1h**). For instance, with 240 nM of PAR28-30 (concentration of PAR chains, calculated for an average of 29 ADP-ribose units from the pool containing PAR with 28, 29, and 30 units) interacting with 100 pM nucleosomes, the resulting decompaction rate is 0.42 s⁻¹.

### Threshold effect in PAR-mediated nucleosome decompaction kinetics: critical length and ionic strength dependence

To assess the impact of PAR length on the nucleosome decompaction process, we first measured the decompaction kinetics of nucleosomes by PAR of different lengths, with 10, 11-12, 14-17, 20-22, 28-30, 35-39, 40-47, and 48-55 ADP-ribose units. Adjacent lengths of PAR were pooled to obtain enough material for the measurements. The observed decompaction rates exhibit some dependence on the PAR concentration but are largely independent of the PAR chain length across this length range (**Figure 2a**). However, when we tested the effect of PAR7 on the nucleosome in the microfluidic mixer, we did not observe decompaction over the 60-second time window accessible in the device (**Figure S4a**, **b**, **c**). We thus switched to a free-diffusion measurement combined with manual mixing and found that the decompaction rate was ∼100-fold slower than for longer PAR chains (**Figure 2a**, **Figure S4b**). This slow decompaction is also observed with PAR lengths of 5, 7, 8, and 9. However, the fraction of decompacted nucleosomes at equilibrium depends on PAR length: the longer the PAR chains, the higher the fraction of decompacted nucleosomes (**Figure 2b**). We thus observe a surprisingly abrupt transition of the decompaction kinetics between short (≤ 9 units) and long PAR (≥ 10 units). Since it has been reported that histones undergo mono-ADP-ribosylation in DNA repair^46–48^, we also tested the effect of mono-ADP-ribose on nucleosome conformation. However, even at a concentration of 400 µM, we did not observe a change in nucleosome conformation (**Figure S5**), indicating that a higher degree of polymerization is essential for the effect of PAR on nucleosome structure.

**Figure 2:**
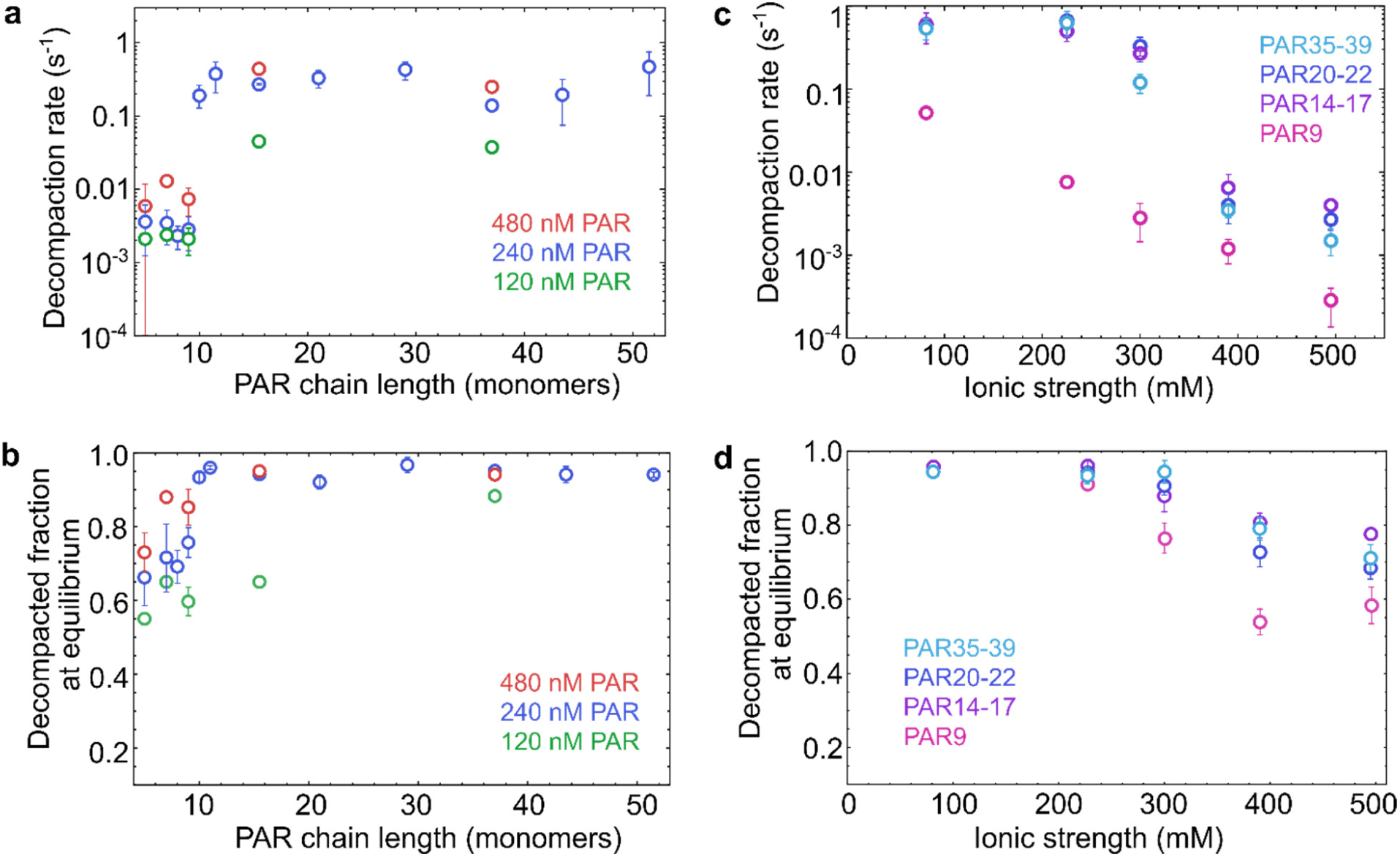
PAR-mediated nucleosome decompaction as a function of PAR chain length and ionic strength. **a**, The apparent decompaction rates of nucleosomes induced by PAR as a function of PAR chain length at three different PAR chain concentrations (ionic strength 300 mM). PAR chains longer than 9 units decompact nucleosomes ∼100 times faster than shorter chains. **b**, The fractions of decompacted nucleosomes at equilibrium against PAR chain length at three chain concentrations (ionic strength 300 mM). PAR chain lengths of 11-12, 14-17, 20-22, 28-30, 35-39, 40-47, and 48-55 were plotted as the average values on the abscissa in **a** and **b**. **c**, Nucleosome decompaction rates induced by PAR of lengths 9, 14-17, 20-22, and 35-39 as a function of ionic strength at a PAR chain concentration of 240 nM. **d**, Fractions of decompacted nucleosomes at equilibrium induced by PAR of lengths 9, 14-17, 20-22, and 35-39 as a function of ionic strength at a PAR chain concentration of 240 nM. All error bars represent standard deviations estimated from at least two repetitions of the measurements.

To assess the role of salt concentration in the process, we examined the kinetics of nucleosome decompaction at various ionic strengths. We chose PAR lengths of 9, 14-17, 20-22, and 35-39 and measured decompaction kinetics at different KCl concentrations. The pronounced salt dependence of the decompaction rates and the fraction of decompacted nucleosomes at equilibrium (**Figure 2c, d**) highlight the role of electrostatic interactions, probably between the negatively charged PAR and the positively charged histone tails^49, 50^. For PAR with 10 or more units, the decompaction rate was largely independent of ionic strength below 300 mM, but it dropped sharply at higher ionic strengths. In contrast, PAR9 exhibited a more gradual reduction in decompaction rate with increasing ionic strength. This transition at ∼300 mM aligns with the previously reported shift of PAR from an extended to a compact state as ionic strength increases^51, 52^, reflecting the overall screening of charge interactions in this ionic strength range.

### PAR affects nucleosome stability via reversible opening and irreversible disassembly

What is the nature of the nucleosome conformations upon PAR binding? The low-⟨*E*⟩ population (0.053±0.002) of nucleosomes formed upon interaction with PAR is close to the ⟨*E*⟩ of 0.035±0.001 of free 197 bp Widom DNA (**Figure 1b, c, d**), and subpopulation-specific fluorescence correlation spectroscopy of the two populations as well as of free Widom DNA shows no significant differences in translational diffusion time (**Figure S1b**). From these results alone, we can thus not decide whether decompaction involves partial DNA opening or complete histone displacement. A different approach is required.

Complete displacement of the core histones is expected to be irreversible at the picomolar nucleosome concentrations we use. We thus first tested the reversibility of the PAR-induced conformational change by digesting PAR with poly(ADP-ribose) glycohydrolase (PARG) to see whether the nucleosomes would recover to the high-⟨*E*⟩ population. To identify suitable concentrations of PARG that completely digest PAR20-22 to monomers, we incubated three concentrations of PARG (0.2 µM, 0.8 µM, and 1.6 µM) with 240 nM PAR20-22 (chain concentration) at 300 mM ionic strength for two hours, respectively, and added the digestion mixtures to the nucleosomes. We did not observe nucleosome decompaction for 1.6 µM and 0.8 µM PARG, indicating digestion of PAR to monomers by PARG (see two upper dashed lines in **Figure 3a**, **Figure S6:** HPLC shows 723 nM PAR20-22 chains were digested to monomers by 1.6 µM PARG after 2 h). The fraction of the nucleosome population with high ⟨*E*⟩ decreased to 64% when the PARG concentration was 0.2 µM (shown as the lowest dashed line in **Figure 3a**), indicating that PARG at this concentration only partially digested PAR, and the remaining PAR led to some degree of nucleosome decompaction. When we added 0.8 µM or 1.6 µM PARG to nucleosomes previously decompacted by PAR20-22, we observed a recovery of compact nucleosomes to around 70% (**Figure 3a**). This observation indicates that most of the decompacted nucleosomes are in a conformation that can be recovered once PAR is removed. However, some nucleosomes cannot be recovered, as would be expected for partial or complete loss of histones from DNA. The fraction of compact nucleosomes recovered decreased with the initial PAR20-22 concentration (**Figure 3b**), suggesting that high concentrations of PAR favor irreversible decompaction of the nucleosome.

**Figure 3:**
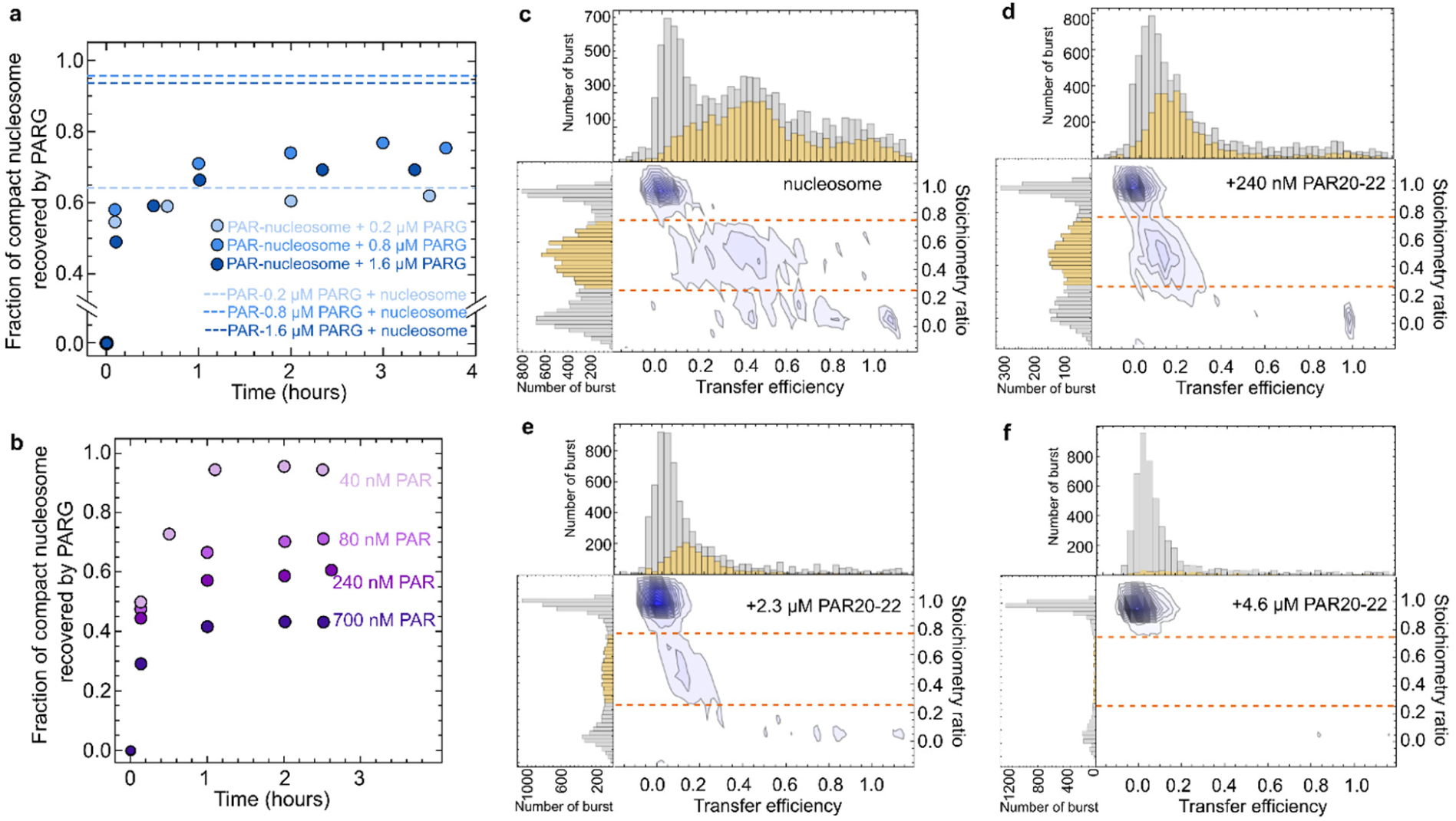
Nucleosome conformational changes via reversible opening and irreversible disassembly by PAR. **a**, Recovery of compact nucleosomes after adding various concentrations of PARG to the PAR-nucleosome complex (240 nM PAR20-22). The dashed lines show the fractions of folded nucleosomes at equilibrium after adding 240 nM PAR20-22 pre-digested by different concentrations of PARG, indicating the extent of digestion of PAR by PARG. **b**, Recovery of nucleosomes decompacted by different concentrations of PAR20-22 after adding 1.6 µM PARG. (The product of 700 nM PAR20-22 incubated with 1.6 µM PARG for 2 hours did not decompact nucleosomes.) **c-f,** Stoichiometry ratio vs. transfer efficiency histograms of EpiDyne®-FRET Nucleosomes labeled with Cy3 (donor) at the 5’ of the 212bp Widom DNA and Cy5 (acceptor) at position 120 of histone H2A, **c**, in the absence and **d**, in the presence of 240 nM PAR20-22 chains (300 mM ionic strength). In the absence of PAR, a donor-only peak (*S* ≈ 1, 〈*E*〉 ≈ 0) and a FRET peak (*S* ≈ 0.5, 〈*E*〉 ≈ 0.4) are visible. Upon addition of PAR20-22, the FRET peak shifted to 〈*E*〉 ≈ 0.15 and the donor-only peak increased in amplitude, indicating that some of the donor-labeled DNA remained associated with the acceptor-labeled histone. In **e** and **f**, 2.3 and 4.6 µM PAR20-22 (chain concentration) were added. The disappearance of the FRET population indicated histone H2A dissociating from the DNA. The dashed lines in c-f indicate the stoichiometry selection 0.25 < *S* < 0.75 for the FRET populations shown in yellow in the transfer efficiency histograms (gray: before cut). All experiments were performed at an ionic strength of 300 mM.

To further test whether nucleosome disassembly involves DNA dissociation from the histone octamer, we performed an intermolecular FRET experiment with commercially available nucleosomes labeled with Cy3 as a donor at the 5’ end of a 212 bp Widom DNA and Cy5 as an acceptor at position 120 of core histone H2A. When 240 nM of PAR20-22 was added, the transfer efficiency shows a shift from high ⟨*E*⟩ to low ⟨*E*⟩, but the stoichiometry ratio stays near 0.5 (**Figure 3c, d**), indicating that the DNA opens up but largely remains bound to the octamer. However, if the PAR chain concentration was increased to several micromolar, the FRET-active population at low ⟨*E*⟩ disappeared, and merely donor-only and acceptor-only populations remained (**Figure 3e, f**), suggesting that H2A dissociated from the nucleosome. Previous studies of nucleosome conformational transitions show that the step-wise disassembly of nucleosomes progresses via the initial loss of H2A–H2B dimers, and eventually DNA fully dissociates from the histones^34–36, 53^.

### Simulations suggest the importance of PAR-histone tail interactions for nucleosome stability

To characterize the conformational ensembles of the compact and decompacted nucleosomes and to identify a possible molecular mechanism by which PAR causes the behavior observed experimentally, we employed coarse-grained molecular dynamics simulations. The DNA was modeled using the 3SPN2.C model^54^, and the histone core (folded segments of the histone octamer) using the AICG2+ model^55^ based on the experimentally resolved crystal structure (1KX5); the disordered histone tails are modeled using a knowledge-based bonded potential^56^. The interactions between the DNA and the histone core are modeled using hydrogen-bond-like interactions with a tunable interaction term^56^, and electrostatic interactions include a screening term to mimic the experimental ionic strengths. The FRET dyes were modeled explicitly^57^ and the FRET efficiencies calculated based on the dye-dye distances to compare with experiment.

The interaction between DNA and histone core was adjusted to reproduce the measured ionic strength-dependent ⟨*E*⟩ of labeled nucleosomes (**Figure 4a** and **Figure S2a**; R = 0.88). The compact nucleosome ensemble observed in the resulting simulations is consistent with previous single-molecule measurements^58^ and cryogenic electron microscopy densities^59^ (**Figure S7**) and corresponds to the DNA fully wrapped around the histone core (**Figure 4c**). The change in linker DNA conformations with ionic strength, probed by the angle of linker DNA with the dyad axis in the plane parallel (α) and perpendicular (β) to the nucleosome disc (**Figure S7c**), reveals the subtle balance between linker DNA repulsion and histone tail-DNA attraction (**Figure S8a**). At low ionic strength, strong linker DNA repulsion and DNA-tail attraction result in larger linker DNA angles, and at high ionic strength, the opposite occurs, explaining the increase in ⟨*E*⟩ observed in the experiments (**Figure 4a** and **Figure S7c**).

**Figure 4:**
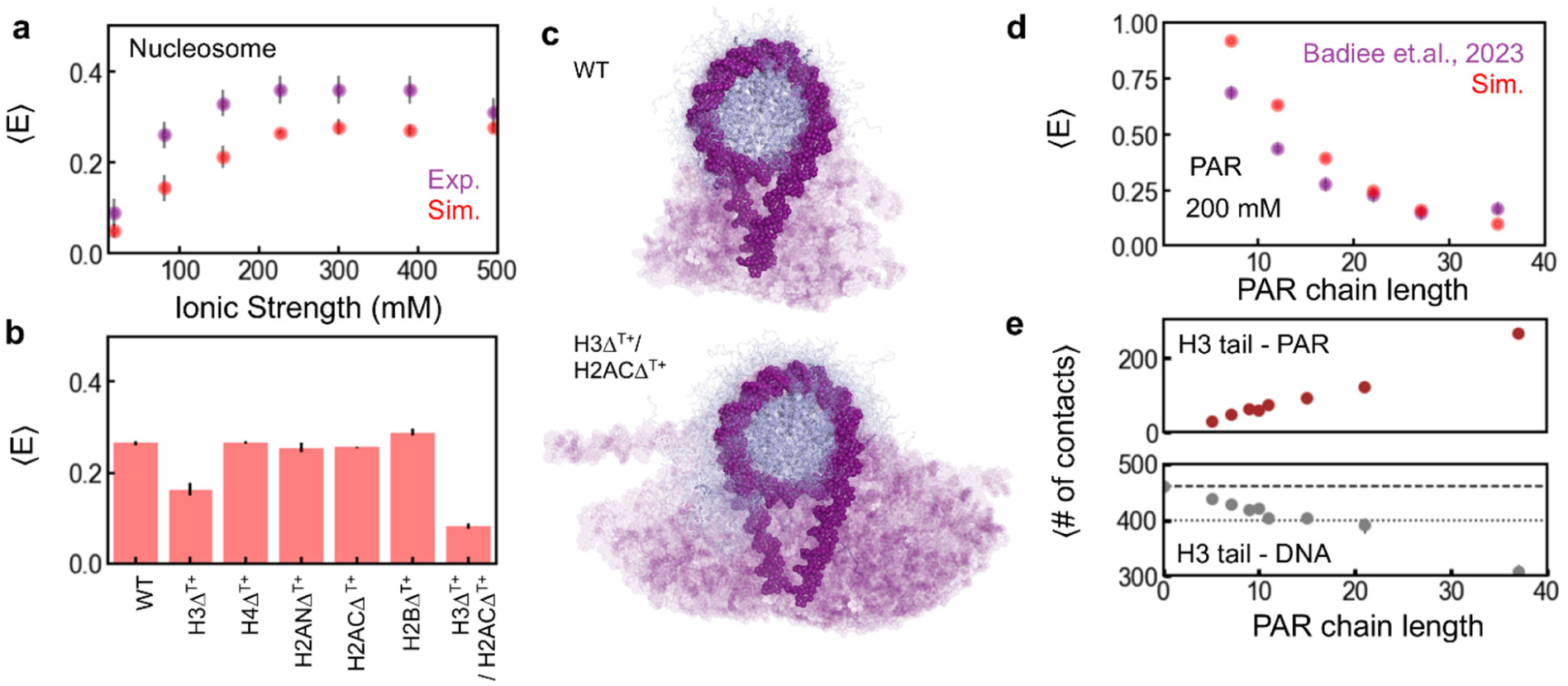
Increasing PAR-histone tail interactions are associated with changing nucleosome conformations in coarse-grained simulations. **a**, The nucleosome model reproduces the trends of the experimentally observed FRET efficiencies of the compact nucleosome as a function of ionic strength. **b**, Mean FRET efficiency, at 300 mM ionic strength, of WT nucleosome and constructs mimicking the loss in histone tail-DNA interactions by mutating all positively charged residues to alanine in IDRs of H3 (H3Δ^T+^), H4 (H4Δ^T+^), H2A (H2ANΔ^T+^ and H2ACΔ^T+^ are N- and C-terminal IDR, respectively), H2B (H2BΔ^T+^) and of both H3 and H2AC (H3Δ^T+^/H2ACΔ^T+^). **c**, Graphical representation of the ensembles of compact (WT) and decompacted (H3Δ^T+^/H2ACΔ^T+^) nucleosome, highlighting the changing linker DNA configurations associated with the loss in contacts of the H3 and H2AC disordered tails with DNA. **d**, The coarse-grained model of PAR approximates the transfer efficiencies of terminally labeled PAR as a function of chain length as probed by single-molecule FRET experiments^51^. **e**, The number of contacts between PAR and H3-tail increases with increasing PAR length and correspondingly disrupts the DNA-histone tail interactions. The dashed and dotted horizontal lines correspond to the average number of contacts in the absence of PAR and in the H3Δ^T+^/H2ACΔ^T+^ tail variant, respectively.

Considering the high stability of the nucleosomes (**Figure S2c**) and the seconds to minutes timescales of PAR-induced unwrapping, we do not expect to be able to observe spontaneous nucleosome unwrapping or dissociation in the simulations. The intrinsically disordered histone tails are highly positively charged and interact with the DNA electrostatically (**Figure S8a**), a key contribution to the stability and dynamics of the nucleosome and its interactions with other proteins^6, 60, 61^. We thus probed the effect of the loss in disordered histone tail interactions on nucleosome conformations by systematically mutating all positively charged residues of the tails to alanine (Δ^T+^); the results suggest that nucleosome unwrapping is dominated by the electrostatic interactions of the H3 and H2A C-terminal tails with DNA (**Figure 4b** and **Figure S8a, b**). Loss of individual histone tail interactions with DNA does not affect linker DNA compaction, except for the loss in H3 tail-DNA interactions (H3Δ^T+^, **Figure 4b, Figure S8b** and **Figure S9**). Loss in DNA-H2A C-terminal tail interactions alone does not favor DNA unwrapping (H2ACΔ^T+^, **Figure 4b** and **Figure S8b**), but the combined effect of H3Δ^T+^/H2ACΔ^T+^ (**Figure 4b, c, Figure S8b,** and **Figure S9**) enhances nucleosome decompaction to a greater extent than H3Δ^T+^ alone. H3Δ^T+^/H2ACΔ^T+^ reproduces the FRET efficiencies of the decompacted state, suggesting that at least 50bp of DNA is unwrapped on either side of the 197bp nucleosome (**Figure 4b** and **c**). DNA unwrapping is primarily due to the loss of DNA-histone tail interactions, as DNA-histone core interactions remain unchanged, with subtle differences at the ends of the nucleosomal DNA (**Figure S8c**). The loss of additional DNA-histone tail interactions (H2BΔ^T+^, H2ANΔ^T+^, H4Δ^T+^), beyond those of H3Δ^T+^/H2ACΔ^T+^, does not affect the linker DNA conformations and nucleosome decompaction, and the corresponding ⟨*E*⟩ does not decrease further and remains consistent with the experimental value (**Figure S9**).

PAR was modeled at a resolution of 3 beads per ADP-Ribose unit, with a charge of -2 on the phosphate beads to represent the diphosphate backbone. FRET efficiencies based on this simplified PAR model are close to previous experiments^51^ (**Figure 4d**), suggesting a reasonable stiffness and charge density of the chains. Since we observed a concentration-dependent effect of PAR in our experiments (**Figure 2**), we simulated nucleosomes in the presence of 1, 25, and 50 PAR chains of varying lengths, ranging from 5 to 37, in the simulation box. We observed that PAR interacts primarily with the histone tails, and the number of contacts increases with both increasing concentration and chain length (**Figure S10**). At low PAR-to-nucleosome ratios, the histone tail-PAR interactions preferentially perturb the interactions of the H4 and H2A C-terminal tails with DNA (**Figure S11**). This finding is in line with previous observations that H4 tails interact with the acidic patch of neighboring nucleosomes, suggesting a weaker intra-molecular interaction with the nucleosomal DNA^62–65^.

At sufficiently high PAR-to-nucleosome ratios, the interactions of all histone tails with DNA are affected. Interestingly, the H3 tail - DNA interactions are severely perturbed by PAR, to an extent equivalent to or greater than by the H3Δ^T+^/H2ACΔ^T+^ mutations for PAR chains longer than 10 units (**Figure 4e**). The loss in H3 and H2A C-terminal tail-DNA contacts with increasing PAR chain length correlates with the change in the decompacted nucleosome population observed in the experiments (**Figure 2b, 4e** and **Figure S11**). H2B tail interactions are perturbed only by longer PAR chains (37mers) at high concentrations (**Figure S11**). In the decompacted state, H2B tails act as the secondary gate keeper, preventing the further unwrapping of DNA through interactions at ±50 bp of the nucleosomal DNA (**Figure S8a**). The loss in H2B tail interactions further disrupts the H3 tail-DNA interactions, destabilizing the nucleosome (**Figure S11**). This molecular mechanism is consistent with only a partial recovery of the compact population upon digestion of the PAR20-22 (**Figure 3**), possibly due to the disassembly of histone octamer or DNA dissociation^34, 35^. The increase in PAR-histone tail contacts with increasing PAR length is linked to an increased lifetime of PAR-histone tail interactions (**Figure S12**) and strongly increases for PAR chains longer than 10 (**Figure 2a**). Although the transition is not as abrupt as observed experimentally, this similarity suggests a coupling between the PAR length-dependent loss in histone tail-DNA interactions and nucleosome decompaction.

## Discussion

By combining microfluidic single-molecule fluorescence spectroscopy under non-equilibrium conditions with coarse-grained molecular dynamics simulations, we find that poly(ADP-ribose) (PAR) controls nucleosome structure by competing with DNA for the interactions with histone tails in a length- and salt-dependent manner. Long PAR chains (≥ 10 units) rapidly decompact nucleosomes, at low concentrations by competing with H3 and H2A C terminal tails and reversibly opening linker DNA. At higher concentrations, H2B tails are outcompeted by PAR, driving more extensive and irreversible DNA dissociation on either end of a 197 bp nucleosome that might be coupled to dissociation of H2A/H2B dimer, the first component typically released in nucleosome disassembly^34–36, 53^ (**Figure 5**).

**Figure 5:**
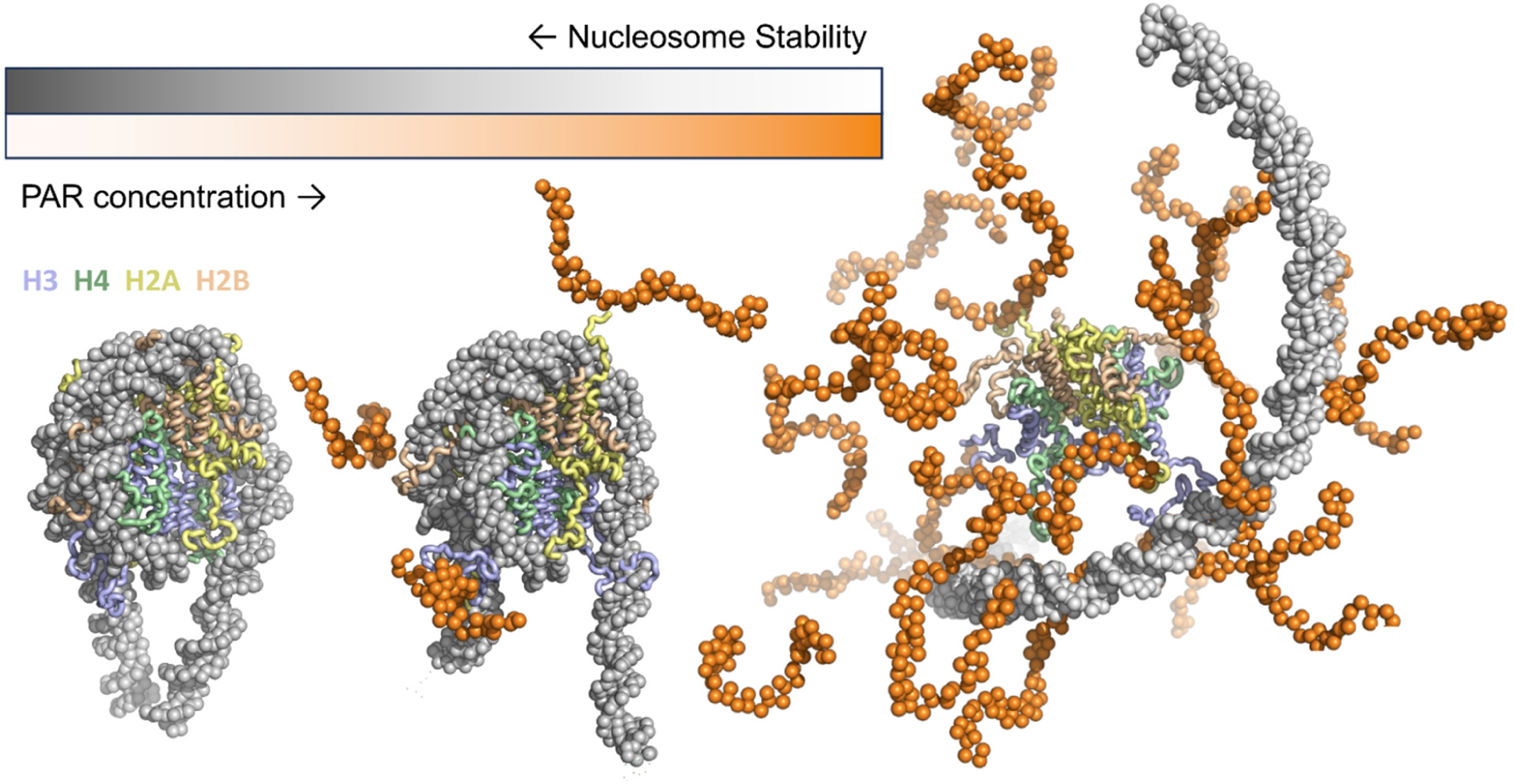
Illustration of the mechanism of changes in nucleosome conformation induced by PAR. In the absence of PAR, the nucleosome assumes a compact configuration, with the linker DNA closed. At low concentrations of PAR, PAR binds to the positively charged disordered tails of histones H3 and H2AC, competing with their interactions with DNA, and resulting in the opening of the linker DNA. At high concentrations of PAR, it displaces multiple histone tails, and the DNA dissociates from the histone octamer.

The sharp kinetic threshold observed experimentally and the drop in DNA–histone tail contacts in the simulations in the range of 10 units suggest a subtle balance in the competition of the interactions between the histone tails, DNA, and PAR. The chain length-dependent nucleosome decompaction is likely to be the result of multivalent interactions: chains long enough to engage several Lys/Arg patches on one or more histone tails bind with much higher effective affinity (**Figure S13**) and have longer dwell times (**Figure S12**), accelerating the transition to the open state. Length-dependent binding of PAR has not only been observed with nucleosomes but is a recurring theme across PAR readers. For instance, the tumor suppressor protein p53 presents much higher affinity for PAR chains longer than 39 units^66, 67^. Similarly, the nucleotide excision repair protein XPA does not interact with short (16-mers) but with long PAR chains (55-mers)^66^. The length-specific interactions of PAR with the readers establish a “PAR code” based on chain length and branching^68, 69^. The inability of mono-ADP-ribose - even at hundreds of micromolar concentrations - to alter the nucleosome conformation provides further support that polymer length and valency, not simply charge, are essential.

At higher ionic strengths, all electrostatic interactions become weaker and this is reflected in the decrease in PAR-mediated decompaction rate at higher salt concentrations (**Figure 2c**). The balance of charge interactions can also have pronounced differential effects on the kinetics: The interactions between PAR chains and the disordered histone tails in the nucleosome are effectively screened above 300 mM ionic strength owing to their high solvent accessibility, leading to a pronounced decrease in the PAR-induced decompaction rate of nucleosomes on the timescale of seconds (**Figure 2c**). The electrostatic interactions between histones and DNA within the densely packed nucleosome, however, are only screened efficiently at higher salt concentrations, resulting in nucleosome disassembly on the timescale of hours at 0.5 M ionic strength (**Figure S2**).

Intracellularly, PAR chains can be covalently attached to and synthesized on multiple targets, including PARP1 and nucleosomal histone tails^18, 19, 70, 71^, in the form of posttranslational modifications as part of the DNA damage response. However, PAR is also synthesized de novo as free polymers from NAD^+^ under such conditions^20^. The two decompaction modes of nucleosomes by PAR we observe may be biologically relevant with respect to these two PAR populations: (i) Transient access via H3/H2AC-tail displacement and linker DNA opening that is readily reversible by PARG would expose DNA around a lesion, increase local breathing, and enhance access for other DNA repair factors, and may be mediated even by free PAR. (ii) The more complete core histone dissociation we observe at high concentrations of PAR above a length of 10 units may resemble primarily the increased local concentrations of covalently linked PAR chains, making DNA more accessible, which may in turn enable recruitment of the complete DNA repair machinery.

## Materials and Methods

### Preparation of fluorescently labeled nucleosomes

#### Preparation of fluorescently labeled oligonucleotides

A ∼10 µl solution of 5–10 nmol oligonucleotide (**Table S1**, thymine modified with a C6-amino linker for the reaction with the succinimidyl ester of the fluorescent dye; Integrated DNA Technologies; Supplementary Table 2) was diluted with 50 µl labeling buffer (0.1 M sodium bicarbonate, pH 8.3). Then 50–100 µg fluorescent dye succinimidyl ester (Alexa488/Alexa594, Life Technologies) dissolved in dimethylsulfoxide was sonicated for 10 minutes and added to the oligonucleotide solution in labeling buffer and the reaction incubated at room temperature for at least two hours. After ethanol precipitation to remove excess dye, the pelleted oligonucleotide was redissolved in 100µl, 95% A (95% 0.1M triethylammonium acetate, 5% acetonitrile), 5% B (100% acetonitrile) RP-HPLC solvent. For assessing the completion of the reaction, a 1 µl sample was diluted with 50 µl of reversed-phase high-performance liquid chromatography (RP-HPLC) solvent A, followed by RP-HPLC using a C18 column (Reprosil-Pur 200 5µm, 250×4mm, Dr. Maisch) with a gradient of 0–100% RP-HPLC solvent B in 50 min. Labeled oligonucleotides were purified preparatively using the same column and gradient as above, lyophilized and resuspended in double-distilled water (ddH2O) to a final concentration of 2.5 µM and stored at -20 °C. The correct molecular weights of the labeled oligonucleotides were confirmed by mass spectrometry.

#### PCR amplification of 197-bp DNA containing the 601 Widom sequence

DNA for nucleosome reconstitution was generated by PCR amplification of a pJ201 plasmid template containing the 197-bp Widom sequence^58^ (gift from Beat Fierz, EPFL), with fluorescently labeled oligonucleotides as primers. The oligonucleotides were designed so that the Widom sequence^37^ was extended by a linker DNA of 25-bp length on either side (See **Table S1**). PCR was typically performed in 10×50 µl volume by mixing in PCR tubes Phusion HF buffer (×1, New England Biolabs), plasmid template (0.02 ng µl^−1^), 0.25 µM forward primer, 0.25 µM reverse primer and deoxynucleotide triphosphates (dNTPs, 0.2 mM each) with ddH2O and 1.0 units of Phusion high-fidelity DNA polymerase (New England Biolabs). Thermocycles included 30 s for initial denaturation at 94 °C, followed by 35 cycles of 20 s for denaturation at 94 °C, 20 s for annealing at 66 °C, and 15 s for extension at 72 °C. The completed PCR reactions were pooled and ethanol-precipitated prior to purification using a Qiagen QIAquick PCR Purification Kit. The concentration of the labeled 197-bp PCR products was determined by UV absorbance. The purity of the 197bp Widom DNA was checked by electrophoresis using 2% Agarose gel stained with GelRed, running at 110V for 30 minutes in 1xTris-acetate-EDTA buffer.

#### Nucleosome reconstitution

Nucleosomes were reconstituted^72^ using 10 pmol of purified DNA containing the 147-bp 601 Widom sequence^37^ flanked by 25-bp linkers. The DNA was mixed with 1.5–1.8 molar equivalents of recombinant core-histone octamer at a final salt concentration of 2 M NaCl on ice. The 30 µl reaction was then transferred to a Slide-A-Lyzer MINI dialysis device (Thermo Fisher Scientific) and dialysed against a linear gradient of buffer with decreasing salt concentration, starting from 10 mM Tris, 0.1 mM EDTA and 2 M KCl, pH 7.5 to 10 mM Tris, 0.1 mM EDTA and 10 mM KCl, pH 7.5, over ∼20 hours. The gradient was created by slowly removing buffer from the dialysis container with a constant flow rate using a peristaltic pump and, simultaneously, supplying fresh buffer with 10 mM KCl using the same flow rate and thus keeping the volume constant. The reactions were then transferred to microcentrifuge tubes and centrifuged for 5 min at 21,000g and 4 °C to remove aggregates; the supernatant was transferred to a new tube. After determining the volumes and the concentrations of the samples (via absorbance at 260 nm), 0.2–0.5 pmol of the reaction products with 25% sucrose were loaded on a 0.7% agarose gel (Invitrogen) and run for 45 min at 90 V with ×0.25 Tris–borate as running buffer. The gels were stained with GelRed (Biotium) for 30 minutes and visualized under UV light.

### Purification of PARP1

Human PARP1 from pQE-TriSystem with a C-terminal 8xHis-tag was purified from Sf21 insect cells. Briefly, three days after infection, cell pellets were resuspended in homogenization buffer (10 mM Tris-HCl pH 7.4, 500 mM NaCl, 10% (v/v) glycerol, 0.1% (v/v) NP-40, 15 mM imidazole, 2 mM β-mercaptoethanol, 0.5 mM phenylmethylsulfonyl fluoride (PMSF), 10 µg/mL pepstatin, 10 µg/mL leupeptin, 10 µg/mL bestatin) and homogenized in a glass homogenizer. The cleared supernatant was incubated for 2 hours at 4°C with ProBond Nickel-Chelating Resin (Thermo Fisher) equilibrated in homogenization buffer. The beads were then washed twice in wash buffer (10 mM Tris-HCl pH 7.4, 10% (v/v) glycerol, 0.2% (v/v) NP-40, 15 mM imidazole, 2 mM β-mercaptoethanol, 0.5 mM PMSF, 10 µg/mL pepstatin, 10 µg/mL leupeptin, 10 µg/mL bestatin) containing 1 M NaCl and twice in wash buffer containing 300 mM NaCl. Bound protein was eluted in elution buffer (10 mM Tris-HCl pH 7.4, 300 mM NaCl, 10% (v/v) glycerol, 0.2% (v/v) NP-40, 250 mM imidazole, 2 mM β-mercaptoethanol, 0.5 mM PMSF, 10 µg/mL pepstatin, 10 µg/mL leupeptin, 10 µg/mL bestatin). The eluted fraction was analyzed by SDS-PAGE and Coomassie staining, aliquoted and shock-frozen in liquid nitrogen. The purification yielded 1.8 mg protein at a concentration of 1.8 µg/µL from 10^7^ insect cells grown adherently in 15-cm dishes.

### Enzymatic synthesis of poly(ADP-ribose)

PAR was produced as a mixture and defined lengths were separated by HPLC. Briefly, 1 mM NAD^+^, 7.5 nM PARP1, 60 µg/mL histone 2a, 50 μg/mL DNA (See **Table S1** for sequence), 100 mM Tris buffer pH 8, 10 mM MgCl_2_, and 1 mM DTT in 1 mL reaction volume were incubated at 37 °C for 15 min. After precipitation of PAR in 20% (v/v) ice-cold trichloroacetic acid, the sample was centrifuged for 10 min at 14 000 g, 4°C. The pellet was washed three times with 1 mL ice-cold absolute ethanol. PAR was released from histone 2a and PARP1 by incubating the washed pellet at 37 °C for 30 min in 0.5 M KOH and 50 mM EDTA. The released PAR was adjusted to pH 7.5 with Tris-HCl buffer (1M Tris-HCl pH 8.0, 37% HCl). DNAse I and MgCl_2_ were added to a final concentration of 0.1 mg/mL and 46 mM, respectively. The mixture was incubated at 37 °C, 550 rpm, for 2 h to degrade the DNA. Then proteinase K and CaCl_2_ were added to a final concentration of 0.2 mg/mL and 0.9 mM, respectively, and incubated at 37 °C, 550 rpm, for at least 2 h to degrade the enzyme 600 μL. Phenol:chloroform:isoamyl alcohol (25:24:1 v/v/v) was then added to the aqueous phase, centrifuged again at 14,000 g for 10 min. PAR in the aqueous phase was finally precipitated by adding 99.8% ethanol to a final concentration of 70% (v/v) and incubated at -20 °C overnight. The precipitate was centrifuged at 9,000 g, 4 °C for 30 min, and the pellet was stored at -20 °C before use. The purified PAR pellet was resuspended in MilliQ water and the concentration in ADP-ribose residues measured by absorbance at 258 nm (extinction coefficient of 13,500 M^-1^cm^-1^).

1 mL of cleaned-up PAR was purified via a DNAPac-PA200 column (Thermo Scientific) using mobile phase A (25 mM Tris buffer pH 9) and mobile phase B (25 mM Tris buffer pH 9 + 1M NaCl) and fractionated at 1 mL/min by the following gradient program: 10 min (0% B), 15min (15% B), 30 min (45% B), 45 min (60% B), 65 min (70% B), 70 min (100% B), 70.01 min (0% B), 80 min (0% B). For each degree of polymerization of PAR, the HPLC elution profiles show two peaks, one corresponding to PAR whose N-terminal phosphoribose was hydrolyzed^73^ and an intact one, respectively (from electrospray ionization mass spectrometry, ESI-MS). The fractions of different lengths of PAR were desalted by ethanol precipitation 3 times and then air dried and stored in - 80°C. The pellets were resuspended in TE buffer (10 mM Tris-HCl and 0.11 mM EDTA, pH8) before use. The different PAR lengths were quantified by ESI-MS.

### Single-molecule fluorescence spectroscopy

Single-molecule experiments were conducted at 22 °C using a MicroTime 200 (PicoQuant) confocal microscope system, including a HydraHarp 400 time-correlated single-photon counting module (PicoQuant). In all kinetic measurements of nucleosome decompaction by PAR, the donor dye was excited using a 485 nm diode laser at 100 µW power (measured at the back aperture of the objective) in continuous-wave mode. Pulsed interleaved excitation^74^ was used to enable alternating excitation of donor and acceptor dyes to quantify FRET efficiencies and stoichiometry ratios of different populations. The donor dye was excited using a 485 nm diode laser at 60 µW power in pulsed mode. For acceptor excitation, the light from a supercontinuum laser (NKT Photonics) operating at 20 MHz repetition rate was passed through a z582/15 band-pass filter (Chroma) and adjusted to an average power of 30 µW at the back aperture of the objective. The sync of the NKT laser triggered the diode laser pulses with a delay of ∼25 ns. Excitation and emission light was focused and collected, respectively, using a high-numerical-aperture water immersion objective (Olympus UplanApo ×60/1.20W). Emitted fluorescence was focused onto a 100 µm pinhole and separated into four detection channels by polarization and donor and acceptor emission wavelengths. Single-photon avalanche diodes were used for detection (Excelitas).

A previously developed droplet-based microfluidic mixer ^33^ was used for measuring the decompaction kinetics from millisecond to second time scale. The microfluidic device was fabricated and operated as described before ^33^. HFE-7500 oil (3M) containing 2% (wt/wt) of the non-ionic tri-block copolymer fluorosurfactant PEG-PFPE2 (RAN Biotechnologies) was used as the carrier phase for all experiments to prevent nucleosome accumulating at the droplet interface. Typically, the flow rate of the continuous phase was set to 6 µL min^−1^, and the flow rate of the aqueous phases was set to 20 nL min^−1^ for measurements, resulting in a droplet velocity of ∼0.6 mm/s in the observation channel, determined by high-speed camera (EoSens 4 CXP, MIKROTRON) recordings during each measurement. The samples in the droplets contained 50–100pM double-labeled nucleosomes with different concentrations of PAR, 140mM β-Mercaptoethanol (Sigma-Aldrich) for photoprotection, and 0.01% (v/v) Tween-20 (Surfact-Amps 20, Thermo Fisher Scientific) for minimizing nucleosome aggregation, in TEK (10 mM Tris, 0.1 mM EDTA, KCl, pH 7.4) buffer with different KCl concentrations (the ionic strength values quoted throughout the manuscript include the 8 mM ionic strength contribution of 10 mM Tris at pH 7.4).

The measurements under equilibrium conditions and of the slow kinetics of nucleosome decompaction induced by short PAR chains (≤9) were performed in µ-Slide sample chambers (Ibidi) following the same sample composition as the kinetic measurements in the microfluidic mixer. Different components were manually mixed in 5 seconds and the measurements started.

For the analysis of the fast nucleosome decompaction kinetics caused by PAR, photon bursts were identified with a threshold of 40 photons per 500-μs bin, and photons from consecutive bins were combined into one fluorescence burst. For the analysis of slow kinetics and the reaction at equilibrium performed in ibidi wells, a threshold of 60 photons per 1-ms bin was used. For each data point of fast kinetics and equilibrium measurements, at least 3,000 bursts were selected, each originating from an individual molecule passing through the confocal volume. Transfer efficiencies were quantified according to *E* = *n*_A_ /(*n*_A_ + *n*_D_), where *n*_D_ and *n*_A_ are the numbers of donor and acceptor photons in each burst, respectively, corrected for background, channel crosstalk, acceptor direct excitation, differences in quantum yields of the dyes, and detection efficiencies^75^. Transfer efficiency histograms from the collected bursts for all time points of a kinetic series were fitted globally with two Gaussian peak functions. Positions of the peaks were shared fit parameters, and the peak widths were constrained to the values expected from shot-noise broadening. For the slow kinetics measured by manual mixing, the reaction was recorded for 20 min and the single-molecule time trace was segmented into one-minute intervals (each containing ∼1000 bursts). For each transfer efficiency histogram, the fraction of compact nucleosome was determined from the fractional area of the high-efficiency peak. The datasets for the decompaction kinetics were fitted globally with a two-state model assuming pseudo-first-order reaction conditions. All data analysis was performed with Fretica, a custom add-on for Mathematica (Wolfram Research) available at https://schuler.bioc.uzh.ch/programs.

### Coarse-grained molecular dynamics simulations

DNA was modeled using the modified 3SPN2.C (three beads per nucleotide) model^54^ with a modified charge of -1.0 on all phosphates, as opposed to -0.6 in the original model, to better reproduce the ionic-strength-dependent trends in experimental measurements. Concurrently, the force constants of the DNA backbone angle and dihedral potential (k_0_= 300 kJ mol^-1^ rad^2^ and k_</_= 6 kJ mol^-1^ rad^2^) were optimized to reproduce the sequence-dependent persistence length of the DNA^54^. Explicit Alexa dyes were modeled as before, matching the experimental dye positions^57^, with three additional linker beads, covalently bonded to the 5’ phosphates, closely matching the chemistry of dye labeling. Proteins were modeled using the AICG2+ model^55^, with each bead representing an amino acid centered at its Cα position. The histone core structure was modeled using the experimentally resolved crystal structure 1KX5^76^ as a reference, and the disordered segments were modeled using a statistical flexible local potential^77^. The interaction between DNA and histone core was modeled using the hydrogen-bond-like interactions with a single tunable interaction energy, as before^56^. The non-specific DNA-histone core interaction energy (5.5 k_B_T) was optimized to reproduce the ionic strength-dependent change in FRET efficiencies observed experimentally (Fig. 4a).

Electrostatic interactions were modeled using a screened Coulomb potential, and the non-electrostatic components were modeled using a short-range solvation potential (hydrophobic scale/HPS model) that included the parameters for explicit dyes^57^. The HPS scaling parameter (λ) for the DNA and PAR beads was set to zero, reflecting their low hydrophobicity. PAR chains were modeled as single-stranded RNA without base stacking and attractive base-pair potentials, with a charge of -2 on the phosphate beads and the phosphate-sugar bond length set to 0.58 nm to mimic the diphosphate backbone. The loss in tail interactions was modeled by *in silico* mutation of positively charged residues (Arg and Lys) to alanine. The nucleosomes, with and without PAR, were placed in the center of a 25×25×25 nm box with periodic boundary conditions. The Langevin dynamics simulations were performed using GENESIS CGDYN^78^ at 300 K with a time step of 20 fs and a friction coefficient γ of = 0.01 ps^-1^.

Mean FRET efficiencies were calculated from the simulations using the inter-dye distances using the Förster equation:

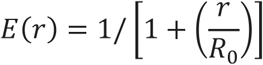

Here, *r* is the distance between the dye beads^57^, and *R_o_*=5.4 nm is the Föster radius of the Alexa dye pair used in the experiments. The values of *E* are compared with the experimental FRET efficiencies. The trajectories were analyzed in Python using the MDAnalysis library^79^, and intermolecular contacts were calculated using a distance cutoff of 1.2 nm. Umbrella-sampling simulations were performed to sample the dissociation of PAR, with a bias applied between the center of mass of the globular domain of the core histones and PAR. For distances between 0 and 16 nm, 16 equally-spaced replicas with a force constant of 10 kJ mol^-1^ nm^-2^ were used. Replica exchange moves between adjacent umbrellas were attempted every 1,000 steps. Potentials of mean force (PMF) were calculated from the equilibrated part of the simulation runs using weighted histogram analysis methods^80^. Molecular visualizations were generated using PyMOL^81^.

## Author information

### Notes

The authors declare no competing financial interest.

## Acknowledgements

We thank Pétur Heidarsson and Lucia Franchini for helpful discussions, and the Functional Genomics Center Zurich for mass spectrometry analysis. This work was supported by the Swiss National Science Foundation (3200-0-239910, 200021_219629 to B.S.), the Novo Nordisk Foundation Challenge grant REPIN – rethinking protein interactions (#NNF18OC0033926 to B.S.), and the Forschungskredit of the University of Zurich (00109396 to S.G.). We used the computational resources of Piz Daint, Alps, and Eiger at the CSCS Swiss National Supercomputing Center. RB was supported by the Intramural Research Program of the National Institute of Diabetes and Digestive and Kidney Diseases (NIDDK) within the National Institutes of Health (NIH). The contributions of the NIH author are considered Works of the United States Government. The findings and conclusions presented in this paper are those of the author(s) and do not necessarily reflect the views of the NIH or the U.S. Department of Health and Human Services.

## Notes

### Competing Interest Statement

The authors have declared no competing interest.

