## Supplementary File for "Time-Resolved Single-Molecule FRET Reveals Length-Dependent Nucleosome Decompaction by Poly(ADP-ribose)"

**Supplementary Information Yang et al.**

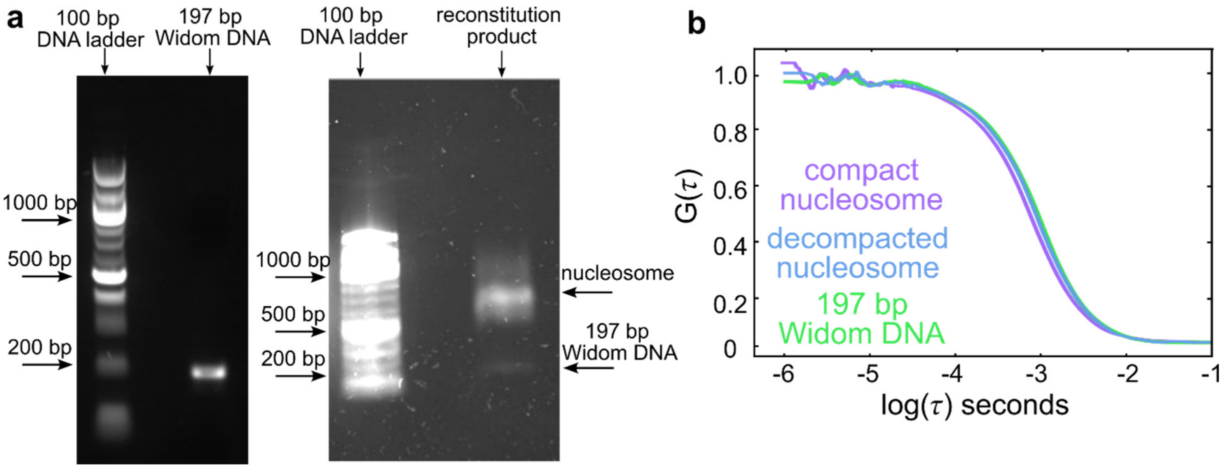


**Figure S1**: Nucleosome reconstitution using 197 bp Widom DNA labeled with Alexa 488 and Alexa 594 at the termini of the linker DNA arms (a1 and b1). **a**, Left: 2% agarose gel of purified 197 bp Widom DNA after PCR stained with GelRed. Right: 0.7% agarose gel of the nucleosome preparation in TEK10 with 25% sucrose stained by GelRed. The bright band at ~700 bp corresponds to intact nucleosomes, and the weak band at ~200 bp indicates the residual presence of free DNA. **b**, Subpopulation-specific fluorescence correlation spectroscopy (FCS) of compact nucleosomes (purple), decompacted nucleosomes (blue), and Widom DNA (green), corresponding to the FRET populations in Fig. 1b-d.

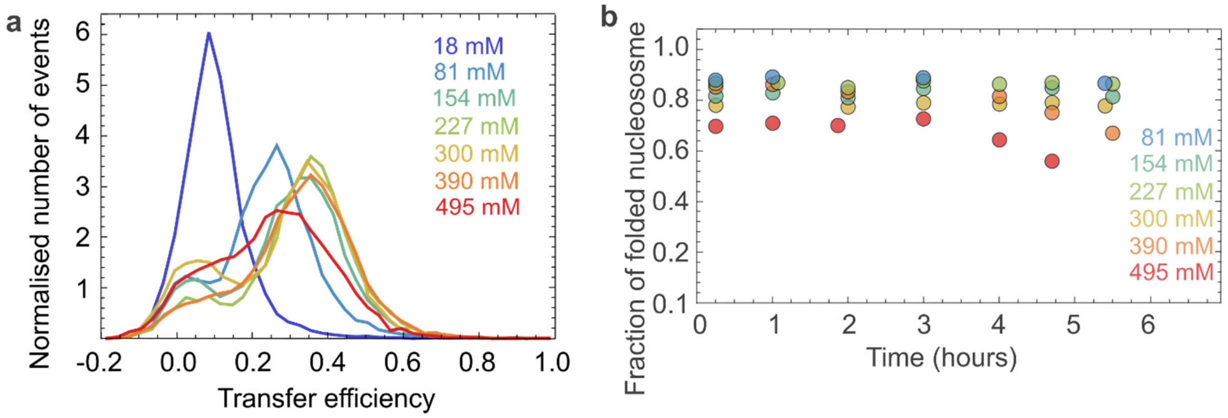


**Figure S2**: **a**, FRET efficiency histograms of nucleosomes terminally labeled at the linker DNA at different ionic strengths. **b**, The fraction of compact nucleosomes observed experimentally over time at different ionic strengths indicates their stability.

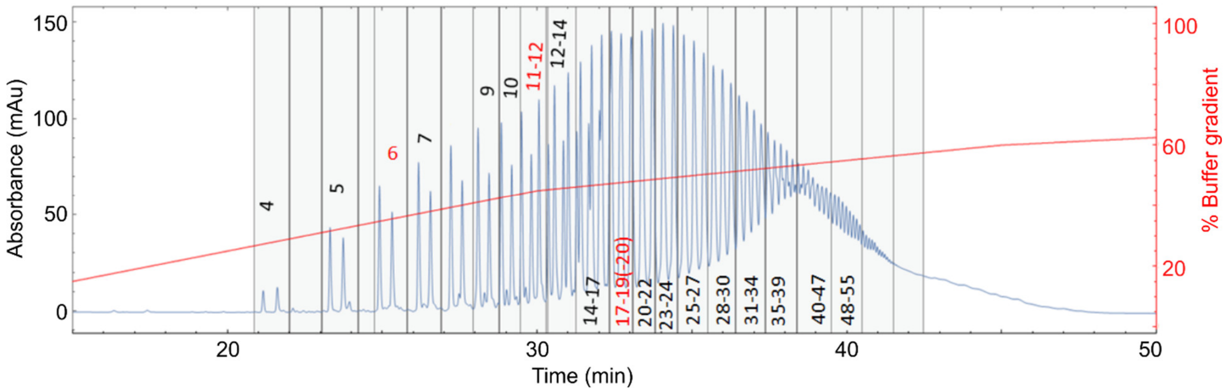


**Figure S3**: HPLC chromatogram of PAR purification after synthesis (absorbance at 258 nm, blue; elution gradient, red) resolves different chain lengths of PAR, with fractions used for measurements indicated by vertical lines. The number at each fraction indicates the number of units. For each length, the first peak is PAR whose N-terminal phosphoribose has been hydrolyzed, and the second peak is unhydrolyzed PAR. The red-labeled fractions were used for electrospray ionization mass-spectrometry for determining the mass of PAR chains and assigning the degrees of polymerization throughout the chromatogram.

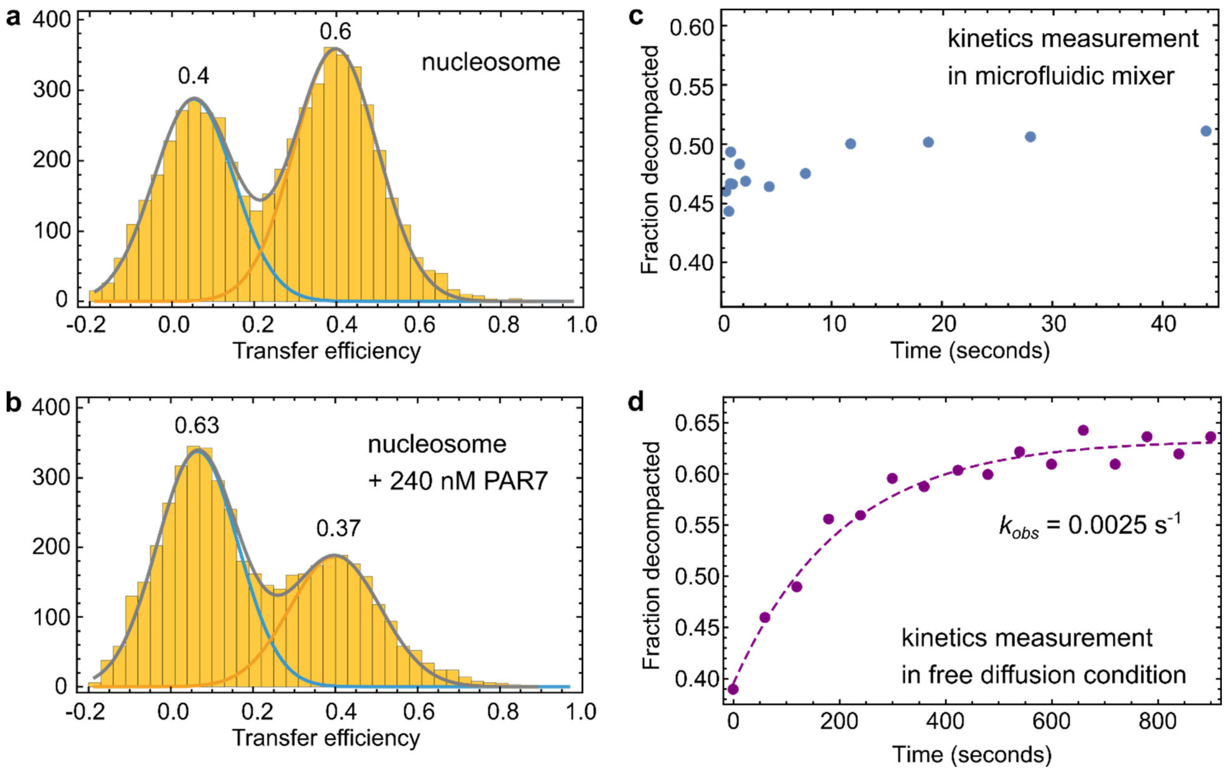


**Figure S4**: **a**, Transfer efficiency histogram of nucleosomes labeled at the linker DNA termini measured in free-diffusion experiments, with a population of compact nucleosomes of ~0.6. **b**, Transfer efficiency histogram of nucleosomes in the presence of 240 nM PAR7 after equilibrium is reached, measured in a free-diffusion experiment. The population of compact nucleosomes is reduced to 37%. **c**, On-chip kinetic measurement of nucleosome-PAR7 interaction from several milliseconds to 44 s. No pronounced change in nucleosome decompaction was observed. **d**, Time series of fraction of decompacted nucleosome upon addition of 240 nM PAR7 measured in free-diffusion experiments and fitted with a two-state model assuming pseudo-first-order reaction conditions (dashed colored line) yields the observed decompaction rate of $k_{\mathrm{obs}} = 0.0025\ \mathrm{s}^{-1}$.

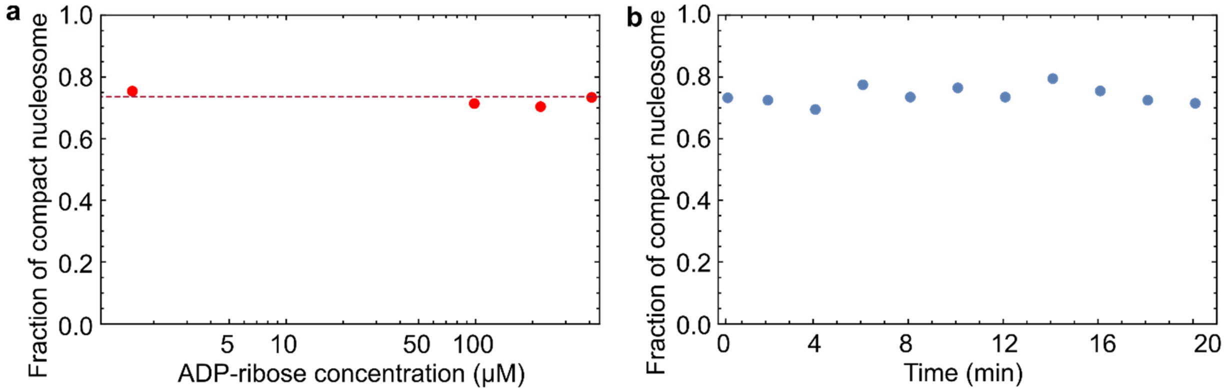


**Figure S5**: **a**, Plot of the fraction of compact nucleosome vs. ADP-ribose concentration (ionic strength 300 mM). Up to 400 μM ADP-ribose, the fraction of compact nucleosomes does not change after 10 mins. **b**, Fraction of compact nucleosome over 20 min with 400 μM of ADP-ribose added.

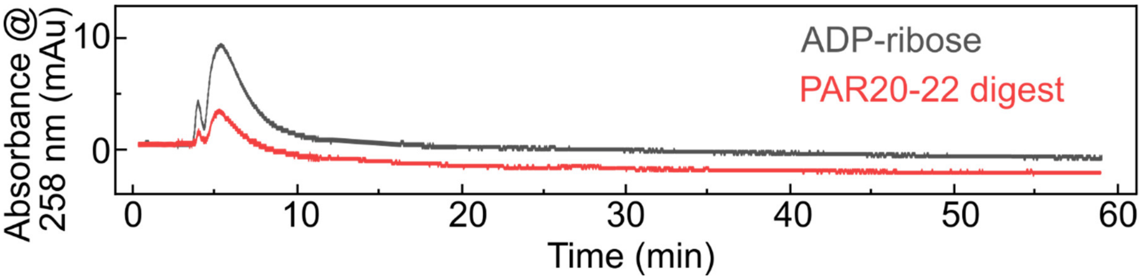


**Figure S6**: HPLC chromatogram of 1.5 nmol ADP-ribose (gray) and 0.5 nmol PAR20-22 (units amount, equivalent to 723 nM PAR chain concentration) digested by 30 pmol PARG (1.6 µM) after 2 h incubation (red). The chromatogram shows that PAR20-22 was digested to monomers by PARG. The mobile phases and elution gradient are the same as for PAR separation by length (Fig. S3).

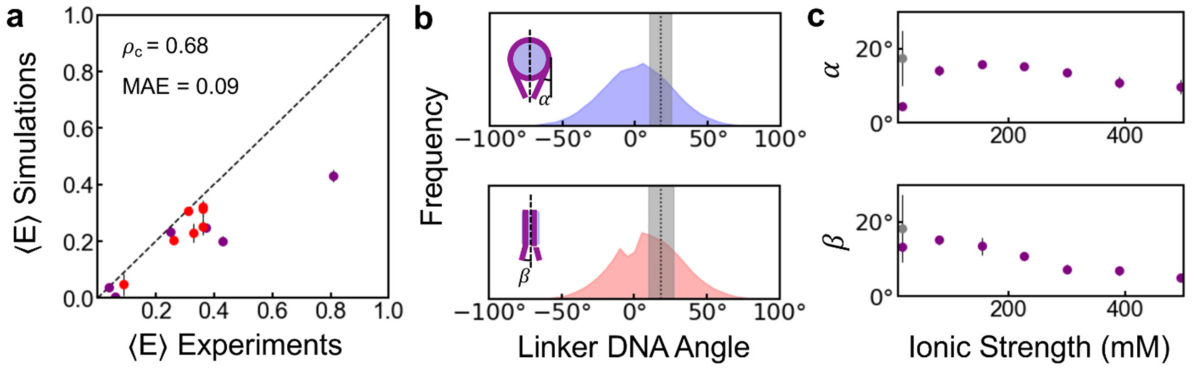


**Figure S7:** **a**, Comparison of average FRET efficiencies from experiment and simulation probed by labeling different DNA positions (purple[1]) and by changing ionic strength (red). The concordance correlation coefficient ($\rho_c$) and mean absolute error (MAE) are shown. **b**, Distribution of linker DNA angles as illustrated in the insets compared against the estimates from cryo-EM data at 18 mM ionic strength (dotted black line; shaded area indicates the standard deviation from the mean[2]). **c**, Linker DNA angles as a function of ionic strength from simulations (purple) and cryo-EM images (grey [3]), and the error bar indicates standard error.

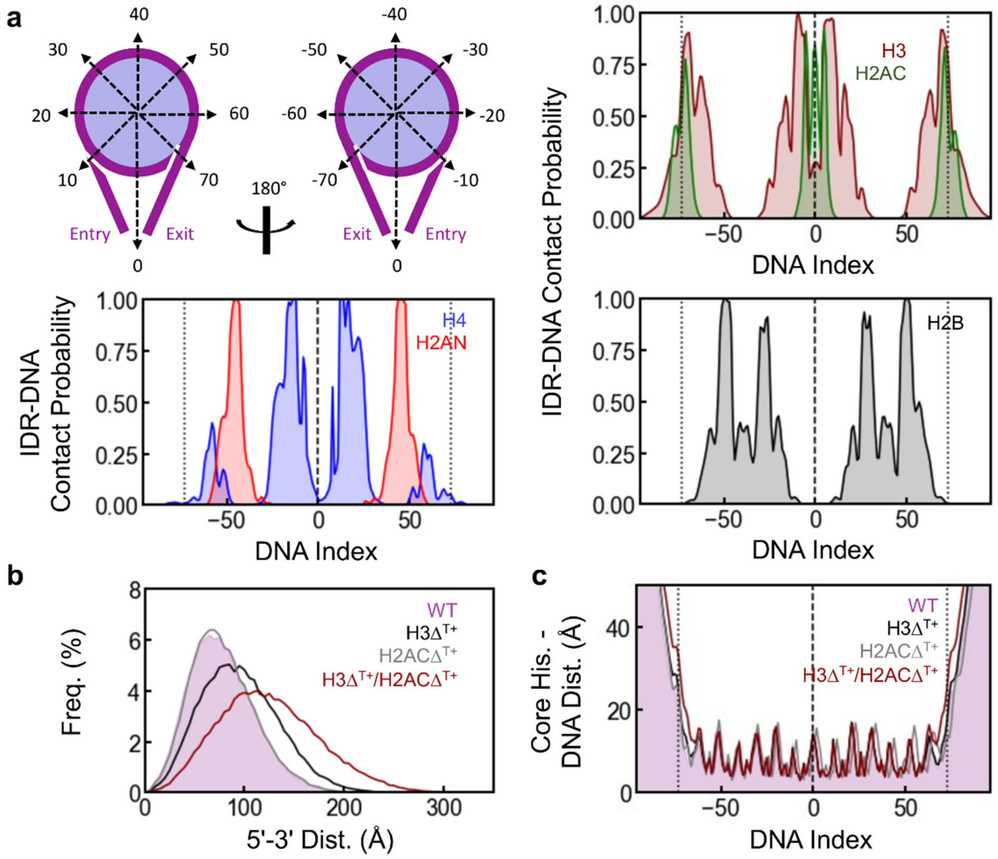


**Figure S8:** Histone-DNA interactions from coarse-grained simulations. **a**, Cartoon representation of the DNA index relative to the nucleosome dyad (index 0). Entry and exit linker DNA are indicated for clarity. Contact probabilities of the five disordered histone tails with DNA at 227 mM ionic strength (different histone tails are shown in different colors as indicated). The vertical dashed and dotted black lines indicate the dyad and nucleosome boundaries (± 73 bp), respectively. **b**, Distribution of the 5'-3' distance of the different nucleosome constructs. **c**, Mean DNA-core histone distance as a function of DNA index. The vertical dashed and dotted black lines indicate the dyad and nucleosome boundaries, respectively. The results suggest that the change in 5'-3' distance observed for $\Delta^{T+}$ variants is primarily due to the loss of electrostatic interactions between histone IDRs and DNA and does not affect the interaction of DNA with the histone core.

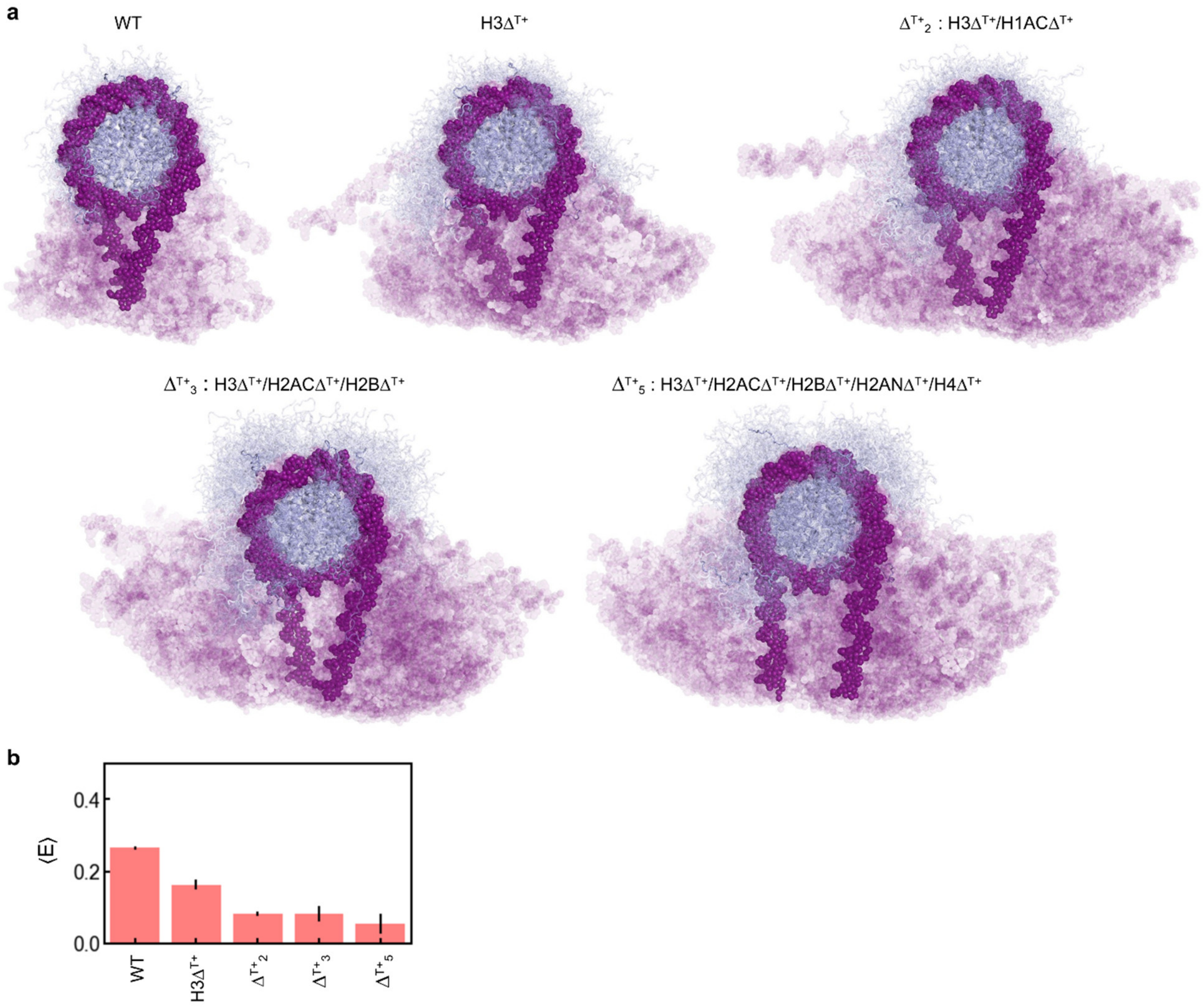


**Figure S9: a,** Conformational ensembles of WT nucleosome and variants mimicking loss in histone tail interactions from simulations (227 mM ionic strength). **b,** FRET efficiency estimated from the simulations for different nucleosome constructs shown in panel **a**. Loss in H3 tail (H3$\Delta^{T+}$) or both H3 and H2AC tail interactions (H3$\Delta^{T+}$/H2AC$\Delta^{T+}$ and $\Delta^{T+}{}_2$ indicate the positive charge neutralizing mutations in two histone IDRs) with DNA increases the distance between linker DNA. Additional removal of tail interactions ($\Delta^{T+}{}_3$ and $\Delta^{T+}{}_5$) does not increase the distance between the linker DNA arms further.

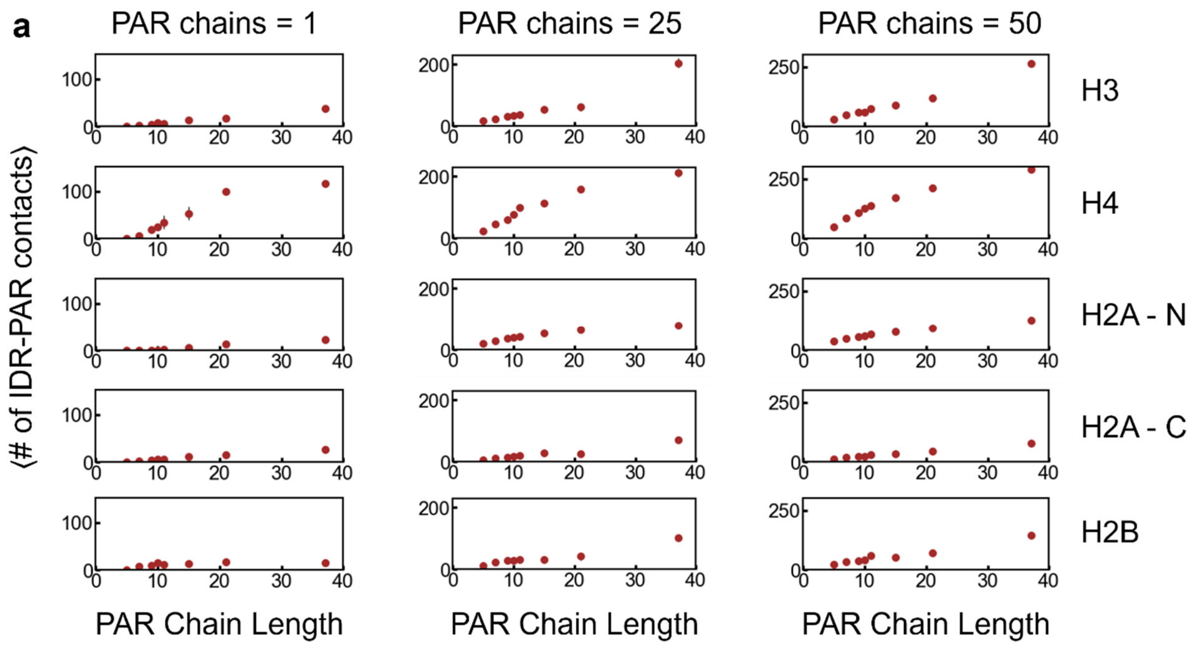


**Figure S10:** Average number of PAR–histone tail contacts with different PAR chain lengths at 227 mM ionic strength as a function of PAR chain length for different numbers of PAR chains per nucleosome included in the simulations. The number of contacts increases with both the length of the PAR chains and the nucleosome-PAR ratio in the simulation box.

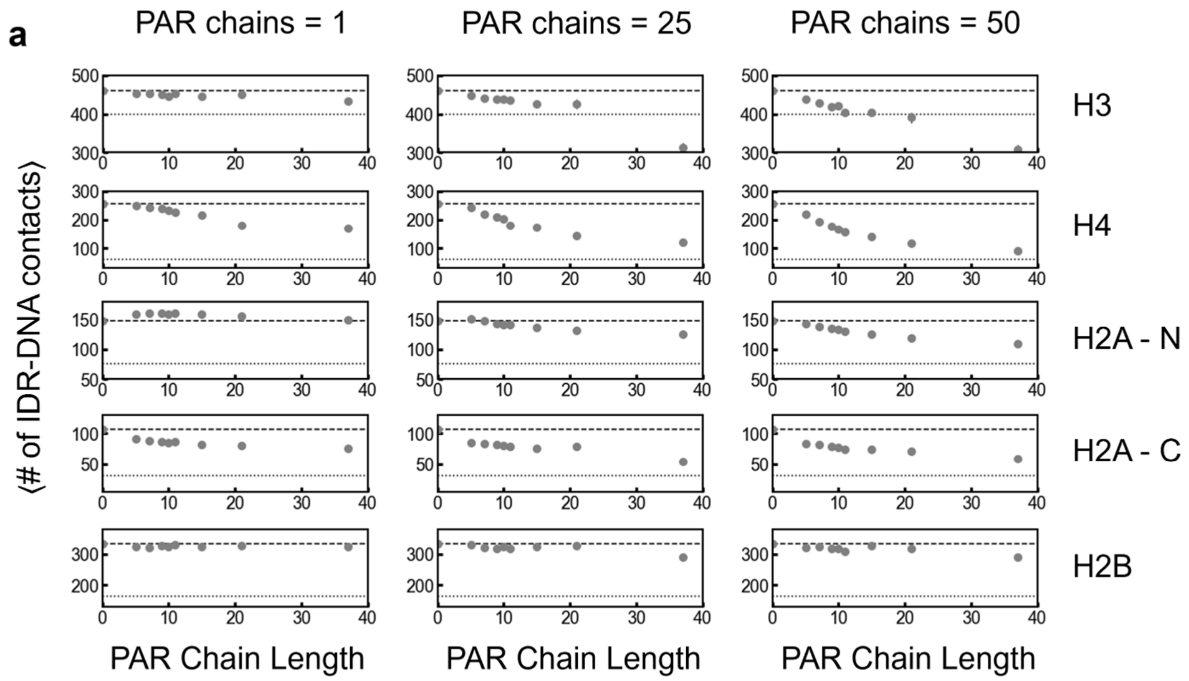


**Figure S11:** Average number of DNA–histone tail contacts in the presence of PAR with varying chain length at 227 mM ionic strength as a function of PAR chain length for different numbers of PAR chains per nucleosome included in the simulations. The number of contacts decreases with increasing PAR chain length and increasing nucleosome-PAR ratio in the simulation box. The dashed horizontal lines indicate the average number of contacts in the absence of PAR, and the dotted horizontal lines indicate the average number of contacts upon mutating all R and K residues in the histone IDRs to A.

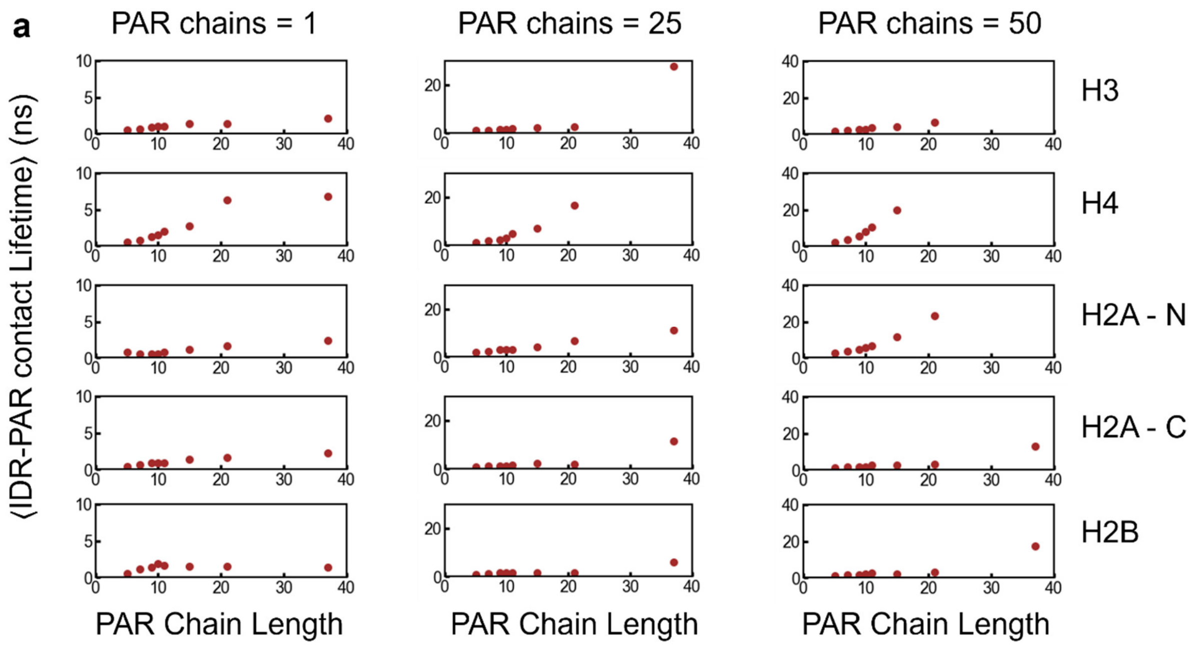


**Figure S12:** Average lifetime of PAR–histone tail contacts with different PAR chain lengths at 227 mM ionic strength conditions as a function of PAR chain length for different numbers of PAR chains per nucleosome included in the simulations. The average contact lifetime increases with both the length of the PAR chain and the nucleosome-PAR ratio in the simulation box, closely resembling the trends of average number of PAR-histone tail contacts in Fig. SF9.

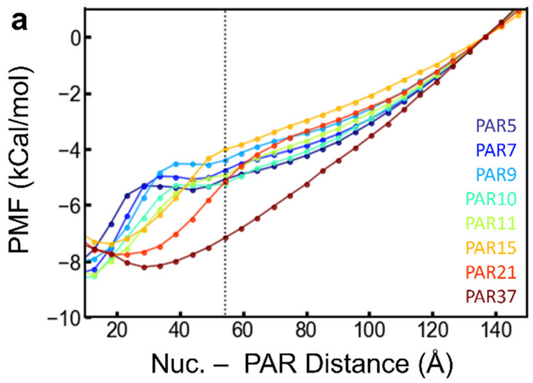


**Figure S13**: Potentials of mean force for unbinding of PAR from the nucleosome from umbrella sampling. The distance was calculated as the distance between the center of mass of the core-histones and PAR, respectively. The dotted line indicates the radius of the nucleosome.

| |
|---|
| 197 bp Widom 601 DNA sequence α-strand:<br>5’-<br>**T**CCATGGACCCTATACGCGGCCGCCCTGGAGAATCCCGGTGCCGAGGCCGCTCAAT<br>TGGTCGTAGACAGCTCTAGCACCGCTTAAACGCACGTACGCGCTGTCCCCCGCGTT<br>TTAACCGCCAAGGGGATTACTCCCTAGTCTCCAGGCACGTGTCAGATATATACATCC<br>TGTGCATGTATTGAACAGCAGTACTAGA-3’ |
| 197 bp Widom 601 DNA sequence β-strand:<br>5’-<br>**T**CTAGTACTGCTGTTCAATACATGCACAGGATGTATATATCTGACACGTGCCTGGAG<br>ACTAGGGAGTAATCCCCTTGGCGGTTAAAACGCGGGGGACAGCGCGTACGTGCGT<br>TTAAGCGGTGCTAGAGCTGTCTACGACCAATTGAGCGGCCTCGGCACCGGGATTCT<br>CCAGGGCGGCCGCGTATAGGGTCCATGGA-3’ |
| Double blunt-ended DNA used in PAR synthesis |
| α-strand: 5’-TGCGACAACGATGAGATTGCCACTACTTGAACCAGTGCGG-3’ |
| β-strand: 5’-CCGCACTGGTTCAAGTAGTGGCAATCTCATCGTTGTCGCA-3’ |

**Table S1**: DNA sequences used in the experiment. Row 1, 2: Sequence of the nucleosome DNA and labeling positions. For DNA labeling, thymine modified with a C6-amino linker was incorporated for the reaction with the succinimidyl ester of the fluorescent dye. Purple labels denote the 3´-end of oligonucleotide primers used to PCR-amplify the corresponding nucleosomal DNA, with previous fluorescence labeling at 5’ ends (labeled red). Row 3-5: Double blunt-ended DNA used in PAR synthesis.
